# Multisensory stimuli improve relative localisation judgments compared to unisensory auditory or visual stimuli

**DOI:** 10.1101/268540

**Authors:** Laura Freeman, Katherine C Wood, Jennifer K Bizley

**Author notes:** equal contribution (Laura Freeman, Katherine C Wood). Corresponding author: Jennifer Bizley.

## Abstract

Observers performed a relative localisation task in which they reported whether the second of two sequentially presented signals occurred to the left or right of the first. Stimuli were detectability-matched auditory, visual, or auditory-visual signals and the goal was to compare changes in performance with eccentricity across modalities. Visual performance was superior to auditory at the midline, but inferior in the periphery, while auditory-visual performance exceeded both at all locations. No such advantage was seen when performance for auditory-only trials was contrasted with trials in which the first stimulus was auditory-visual and the second auditory only.

## INTRODUCTION

Both auditory(Mills, 1958; Makous and Middlebrooks, 1990; Charbonneau et al., 2013; Wood and Bizley, 2015; Carlile et al., 2016) and visual localisation acuity declines with eccentricity (Mateeff and Gourevich, 1984; Perrott et al., 1993; Charbonneau et al., 2013; Carlile et al., 2016). Few previous studies have attempted to directly compare spatial acuity for auditory and visual stimuli throughout the visual field and focus instead on the spatial capture observed when spatially separated auditory-visual signals are presented (Howard and Templeton, 1966; Bertelson and Radeau, 1981). Two exceptions to this are Perrot et al., (1993) and Charbonneau et al (2013). Both determined that both visual and auditory localisation judgments declined as stimuli move from central to peripheral space. However, the studies produced conflicting results, and neither study perceptually matched stimuli across modalities. Perrott et al., did not test bimodal stimuli, and reported equivalent auditory and visual performance, while Charbonneau reported superior visual performance and no advantage for auditory-visual stimuli but on every trial an auditory-visual reference was provided and only the target varied in modality.

The aims of this study was to determine (i) how relative localisation judgments vary throughout frontal space for *equally-detectable* auditory and visual signals and (ii) whether an auditory-visual signal conferred a processing advantage over the most effective unisensory stimulus. Finally, because we observed a clear multisensory benefit, we also included stimuli in which an auditory-visual reference was followed by an auditory only target. It was hypothesised that localisation acuity would decline with eccentricity for both auditory and visual signals but that at central locations (i.e. the fovea) vision should provide the most accurate estimate of source location, whereas at more peripheral locations sound localisation would be more accurate than visual localisation.

## METHODS

### A Participants

This experiment received ethical approval from the UCL Research Ethics Committee (3865/001). 14 normal hearing adults between the ages of 18 and 35 participated in Experiment 1. Two participants were excluded due to poor performance (average d’ <0.5). 9 of the remaining 12 participants participated in Experiment 2. All participants had no reported hearing problems or neurological disorders.

### B Testing chamber

For testing, participants sat in the middle of an anechoic chamber surrounded by a ring speakers arranged at 15° intervals from −67.5° to +67.5° (Figure 1A). Each speaker had a light emitting diode (LED) mounted immediately below it. The participants’ heads were kept in a stationary position and supported there by a chin rest. Participants were asked to maintain fixation on a fixation cross located on the speaker ring at 0° azimuth and their head and eye position were remotely monitored with an infra-red camera.

**Figure 1 (color online).**
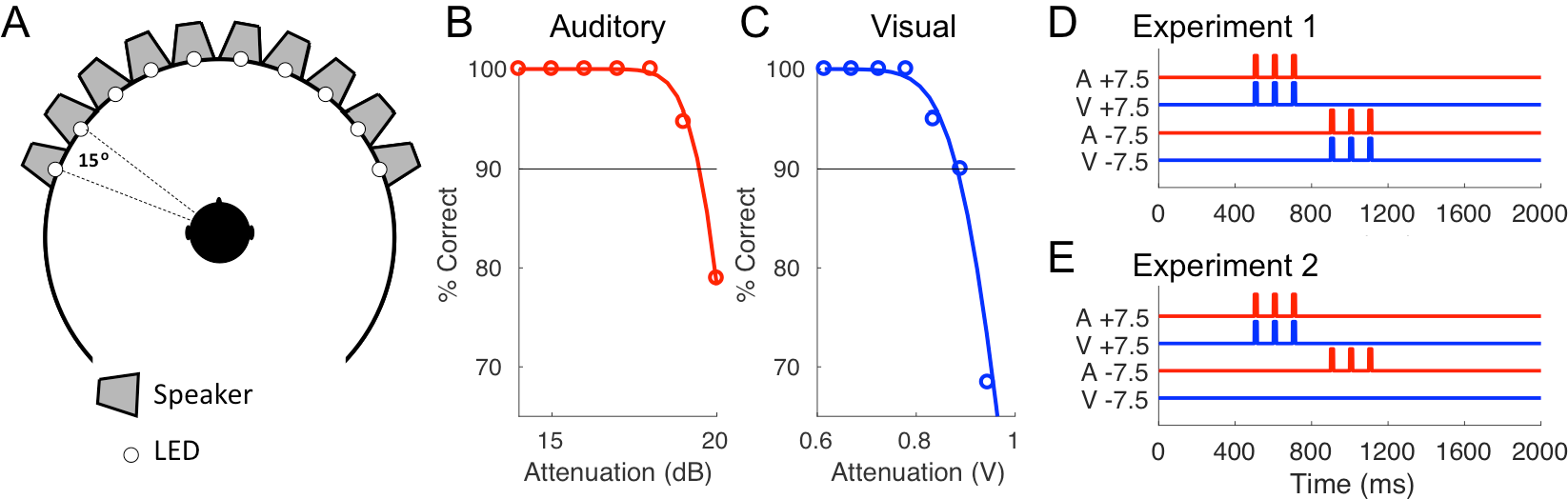
A Schematic of the testing chamber B, C Example threshold function for auditory (B) and visual (C) detection abilities. D, E, schematic of the trial structure for Experiment 1 (D, AV trial) and Experiment 2 (E, AV reference trial) showing for one example trial in which the relative location of the stimulus shifts leftwards from +7.5 degrees to −7.5 degrees.

### C Stimuli

All stimuli were generated in MATLAB and presented using the PsychToolBox extension (Brainard, 1997) at a sampling frequency of 48 kHz. Participants reported the location of a target stimulus (left or right) relative to a preceding reference stimulus. In the Auditory (A) condition 3 pulses of white noise were presented from a reference speaker, followed by 3 pulses of white noise from a target speaker. In the Visual (V) condition three pulses of light were emitted from a reference LED mounted on a speaker, followed by three pulses of light from a target location. In the Auditory-Visual (AV) condition in Experiment 1 spatially and temporally coincident light and sound pulses were presented. In Experiment 2 spatially and temporally coincident sound and lights were presented at the reference location, and only the auditory stimulus was presented at the target location. Auditory stimuli were broadband noise bursts (as in Wood and Bizley, 2015). Reference and target speakers were always separated by 15° (Fig. 1A). Stimulus pulses were 15 ms in duration, cosine ramped with 5ms duration at the onset and offset of each pulse. Pulses were presented at a rate of 10 Hz with a 185 ms delay between the end of the final pulse at the reference speaker and the first pulse at the target speaker in order to aid perceptual segregation of the reference and the target. The pulses were embedded in a noisy background generated by presenting independently generated auditory and visual noise from each speaker/LED. The amplitude was varied every 15 ms with amplitude values drawn from a distribution whose mean and variance could be controlled (see Wood and Bizley, 2015). In these experiments the mean noise level across all speakers was 63 dB SPL (calibrated using a CEL-450 sound level meter) and the signal attenuation was set for each participant by performing a threshold measurement. At the start of each trial the noisy background was ramped on with a linear ramp over 1 second and ramped down over 1 second at the end of the trial. The stimulus pulses, which constituted the reference and target, were presented between 50 and 1000 ms after the noise reached its full level. Stimuli were presented by Canton Plus XS.2 speakers (Computers Unlimited, London) and white LEDs via a MOTU 24 I/O analogue device (MOTU, MA, USA). For auditory stimuli the MOTU output was amplified via 2 Knoll MA1250 amplifiers (Knoll Systems, WA, USA). Testing runs were divided into blocks of trials lasting approximately 5 minutes. At the end of each block the participant could take a break and choose when to initiate the next block. Participants performed 15 trials for each reference location / direction /modality combination.

### D Threshold

In order to perform the auditory and visual task at equivalent levels of difficulty an initial *threshold* test was performed. In this task participants were oriented to face a speaker at the frontal midline (0° azimuth). The reference stimulus was always presented from this speaker/LED, and the target was presented from a speaker/LED at either −60° or +60°. Auditory and visual stimuli were presented in separate testing blocks. Participants reported the direction in which the stimulus moved using the left and right arrows on a keyboard to indicate −60° and +60°, respectively. Auditory stimuli were presented at 10 different SNRs by varying the signal attenuation in 1 dB steps over a 10 dB range, and visual stimuli were presented at 7-10 SNRs by varying voltage values driving the LEDs. Percentage correct lateralisation scores were fit using binomial logistic regression and the threshold value (90% correct) was extracted from the fitted function. The aim was to present stimuli at a level that was clearly audible/visible, but difficult enough to be challenging for the subsequent relative localisation task. The threshold therefore served both to match difficulty across participants (as in Wood and Bizley, 2015) and sensory modalities.

### H Analysis

Overall performance was assessed using signal detection theory to calculate sensitivity index (*d*’) statistics for participants, ability to discriminate whether a target sound moved left or right (Green and Swets, 1966). Performance was estimated across reference-target pairs of the same locations (so that the change in localisation cues for left moving and right moving trials were equivalent) and considered relative to the mean location of that speaker pair. Multisensory gain was calculated as the improvement in performance in the multisensory condition relative to the best unisensory condition (in Experiment 1) or the unisensory auditory stimulus (in Experiment 2). Since performance varied with azimuthal position, values were expressed as a % relative to the best unisensory performance for that eccentricity (Charbonneau et al., 2013). Reaction times were compared to predictions of the race model (Miller, 1982) using methods provided by Ulrich et al., (2007). Group level statistical analysis was performed in SPSS (v24, IBM) using repeated measures analysis of variance (ANOVA). Two-way repeated measures ANOVA were performed to determine the impact of modality and spatial location on sensitivity, bias and reaction time measures. One-way repeated measures ANOVA was used to determine the impact of eccentricity on multisensory gain or location within a modality.

## RESULTS

Before participating in Experiment 1 listeners performed two short detection-in-noise threshold tests. These served to match the detectability of signals across modalities by assessing performance in a reduced version of the task across a range of signal attenuations (Fig 1B,C). This step is critical as it allows us to test each modality at an equivalently difficult level so that we can directly compare localisation ability across auditory and visual signals, it further serves to match difficulty across participants.

### Experiment 1

Experiment 1 tested the ability of listeners to perform relative localisation judgments with auditory (A), visual (V) or spatially and temporally coincident auditory visual (AV) signals, presented at their pre-determined signal attenuations. Performance varied throughout azimuthal space (Fig. 2A) with the best performance being obtained for stimuli close to the midline, and performance dropping off at more lateral locations. V performance, although superior to A at the midline, dropped with eccentricity more dramatically such that A performance was superior in the periphery. AV performance exceeded A and V at all locations except for stimuli crossing the midline, where performance was close to ceiling for both V and AV stimuli. Both stimulus modality (F_(2,22)_ =20.8, p=0.0006) and location (F_(8,88)_ = 24.9, p=1.25e-19) influenced *d’*, with a significant modality x location interaction (F_(16,176)_=20.8, p=1.0934e-9). Pairwise post-hoc comparisons revealed that AV performance was significantly different from both A and V (which were statistically indistinguishable) and that central reference locations were significantly different from peripheral ones (Table 1). Multisensory gain was calculated by comparing *d′* values obtained in the AV condition with those in the best unisensory condition, with data folded across space to determine how eccentricity impacted multisensory gain (Fig 2D). T-tests (Bonferoni corrected for 5 locations) indicated that multisensory gains were non-zero at 15°, 30° and 60° (p<0.01) and gain did not vary significantly with eccentricity (effect of eccentricity on multisensory gain: F_(4,44)_=1.82, p=0.142).

**Table 1.** Post-hoc pairwise comparisons (Bonferoni corrected) for the effect of spatial position in experiment 1.

| Mean Location | -60 | -45 | -30 | -15 | 0 | 15 | 30 | 45 | 60 |
| --- | --- | --- | --- | --- | --- | --- | --- | --- | --- |
| Performance different (p<0.05) at: | -30, -15, 0, 15, 30 | -30, -15, 0, 15 | -60,-45 | -60, -45, 30, 45, 60 | -60, -45, 30, 45, 60 | -60, -45, 15, 60 | -60, -15, 0, 15, 60 | 0 | -15, 0, 15, 30, 45 |

**Figure 2 (color online).**
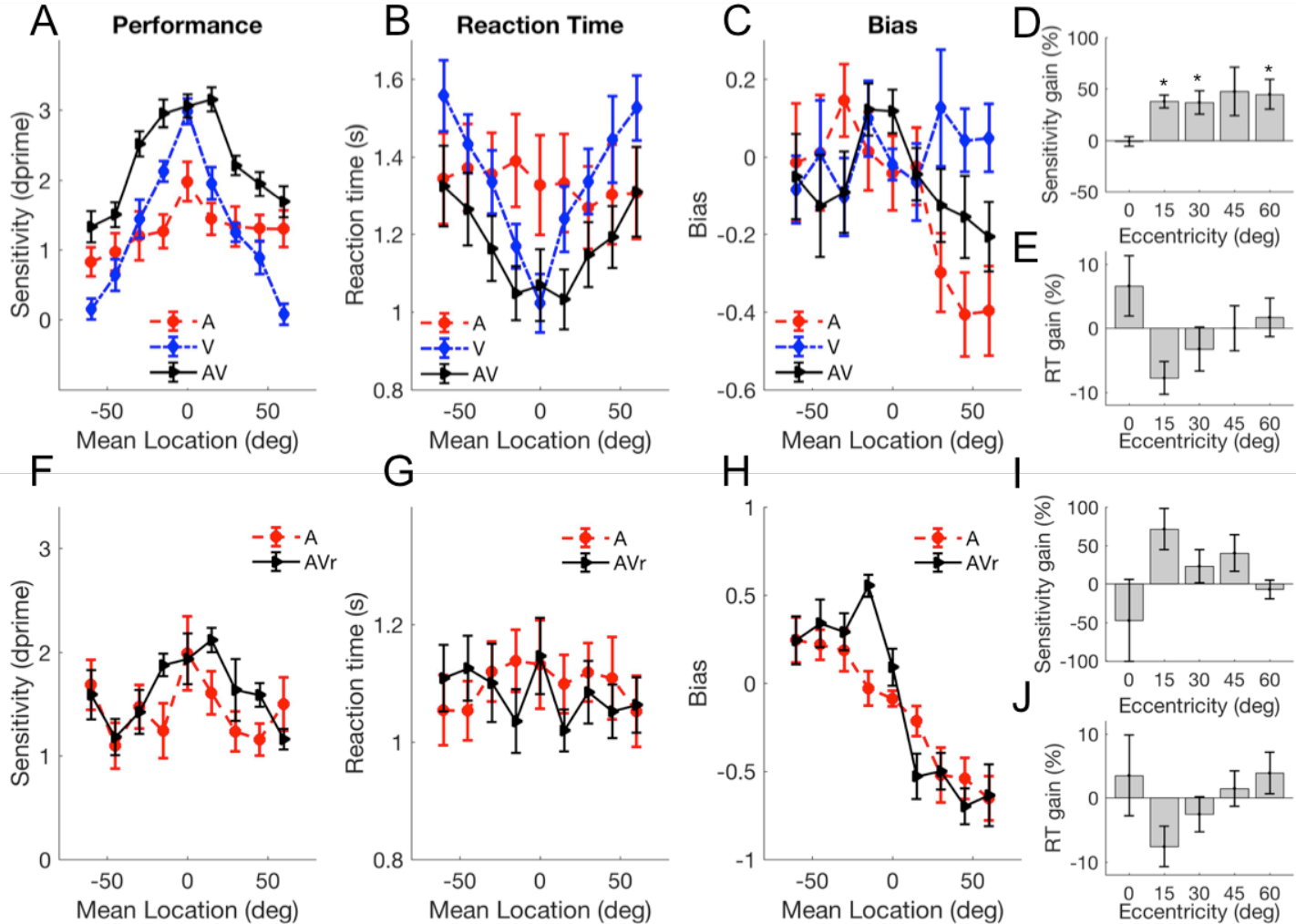
Mean (±SEM) (A) d’ scores for A, V and AV trials as a function of the mean reference-target location, (B) reaction times, (C) bias, (D) sensitivity gain (% gain relative to best unisensory performance), E, reaction time gain (% relative to fastest unisensory). ^*^ indicate values are significantly non-zero (p<0.05 corrected for 5 comparisons). F-J, as A-E, but for Experiment 2.

Reaction time measures (Fig. 2B) for relative localisation judgments with A and V stimuli showed distinct patterns: V reaction times rose monotonically with increasing eccentricity (one way ANOVA of location on V reaction times F_(8,88)_=16.1 p <0.001), while A reaction times were consistent across space (F_(8,88)_=0.85 p=0.57). AV reaction times showed an intermediary pattern of variability increasing more gradually with eccentricity (AV: F_(8,88)_=6.94 p <0.001) and, with the exception of the central location, always being faster than either modality alone. A two-way ANOVA investigating the influence of position and modality on reaction time revealed effects of both location (F_(8,88)_=10.34 p=4.3405e-10) and modality (F_(2,22)_=4.46, p=0.024) with a significant modality x location interaction (F_(16,176)_=5.73 p=6.7686e-10). Post-hoc analysis revealed that AV reaction times were significantly faster than both auditory and visual reaction times. While AV reaction times were significantly faster than either modality alone, they did not violate the race-model (Miller, 1982;), p>0.05 at all locations) and when reaction times were expressed as multisensory gain (Fig. 2D,E), no location had a significantly non-zero gain (t-test against zero, Bonferoni corrected p<0.01).

Bias measures were calculated for performance in each modality (Fig.2C). For both V and AV trials performance was constant across space (one way repeated measures ANOVA, AV: F_(8,88)_=1 27, p=0.270 V: F_(8,88)_=0.64, p=0.742) whereas for A bias was influenced by spatial position (F_(8,88)_=2.92, p=0.006). Consistent with this, a two-way repeated measures ANOVA directly comparing these values revealed no effect of either modality (F_(2,22)_=2.76, p=0.085) or spatial position (F_(8,88)_=1.279, p=0.269), but a significant modality x position interaction (F_(16,176)_=2.23 p=0.006; Fig.2C). In summary, AV stimuli conveyed an advantage in both performance and reaction time compared with the best unisensory stimulus, throughout frontal space.

### Experiment 2

Experiment 2 aimed to determine whether the improvement in relative localisation ability for auditory-visual stimuli could be observed by presenting an AV reference stimulus and an auditory-only target. Nine of the 12 participants from Experiment 1 performed Experiment 2 which included trials which were A-only for both reference and target, and AV reference, A-target trials. An AV reference provided no advantage over an A reference when the target was A alone (Fig 1E): Performance varied weakly with reference location (F_(8,64)_=2.391, p=0.025, post-hoc pairwise comparisons all p>0.05), but not modality (F_(1,8)_=2.56, p=0.148), nor was there a significant modality x location interaction (F_(8,64)_=1.788, p=0.096, Fig. 2F). Reaction times were also uninfluenced by an AV reference stimulus (spatial position; F_(8,64)_=1.06, p=0.5, modality; F_(1,8)_=1.179, p=0.309, Fig. 2G). Consistent with an AV reference offering no perceptual advantage, measures of multisensory gain were not significantly different from zero (t-test, all p>0.05, corrected for 5 comparisons, Fig.2I,J). Finally we considered bias: consistent with auditory performance in Experiment 1, both auditory and AV reference conditions showed very similar patterns of bias, with listeners tending to show positive biases in left space, and negative biases in right space indicating a preference to respond towards the midline (spatial position; F_(8,64)_=16.46, p=0.000, modality; F_(1,8)_=1.179, p=0.309, modality x position interaction F_(8,64)_=3.43 p=0.002; Fig.2H). Thus the multisensory enhancement seen in Experiment 1 required that both stimulus intervals contained a multisensory stimulus.

## DISCUSSION

In these experiments we tested the accuracy with which observers could discriminate 15° shifts in location between sequentially presented reference and target stimuli. Difficulty matched auditory and visual stimuli were used so that performance could be directly compared across modalities. Visual accuracy was highest for central locations and fell off sharply at more peripheral locations. Auditory accuracy was highest at the midline, and also declined at more peripheral locations. However, the change in auditory relative localisation ability with eccentricity was much smaller in magnitude (**Δ** *d* ‘ change of 1.2 for A, compared to **Δ** *d*’ = 2.9 for V) than for visual ability. Performance for auditory-visual stimuli also varied throughout space and, except at the midline where performance matched V (and performance was at or close to ceiling), was better than either A or V at all locations. AV stimuli were processed faster than A or V. Consistent with previous studies V reaction times increased with eccentricity, and AV reaction times mirrored these, whereas processing time was not contingent on eccentricity for A-only stimuli.

These results emphasise that the advantage conferred by visual stimuli exists only in central regions closest to the fovea; at more lateral locations auditory stimuli are more accurately localised but that integrating stimuli offers an advantage throughout space. These findings mirror those of Perrott et al., (1993); although they demonstrated no statistical difference between auditory and visual stimuli, the group data for their 4 observers suggest that visual acuity exceeded that of auditory acuity at 0° (minimum visual angle, MVA=0.5°, minimum auditory angle, MAA=1°), was equivalent (roughly 2°) at 20°, and auditory acuity exceeded visual acuity at more lateral locations (for example at 80 degrees reference MAA=4°, MVA=7°). Charbonneau et al., (2013) performed a similar experiment to the present study, except that they only varied the modality of the target stimulus: a spatially congruent AV reference was presented on every trial. They reported that AV performance matched that of V, and exceeded A, at all locations. The difference in the results presented here and those in Charbonneau et al., (2013) is likely explained by our presenting matched-detectability stimuli across modalities which provided the opportunity to make direct comparisons in spatial acuity.

Where and how multisensory signals are integrated for decision-making is likely to be task and stimulus dependent (Bizley et al., 2016). The improvement in performance observed for multisensory stimuli could arise through multiple mechanisms: it might be that by cueing cross-modal spatial attention to a particular region of space with the reference stimulus performance is enhanced(Spence and Driver, 1997); it may be that early cross-modal integration of auditory and visual signals within auditory cortex (Bizley and King, 2008) enables the visual stimulus to improve the representation of the sound in auditory cortex; alternatively separate auditory and visual estimates of the relative location of the reference and target sound might allow weighted integration at a later decision-making stage (Alais and Burr, 2004). While relating localisation acuity and accuracy is non-trivial (Moore et al., 2008) an improved reference representation should facilitate improved performance: The results of Experiment 2, in which an AV reference stimulus did not enhance the ability of observers to discriminate the direction of a subsequent auditory target, is therefore most consistent with the third option: that the improvement in performance seen for multisensory stimuli results from the integration of separate auditory and visual decisions.

## Acknowledgments

This work was supported by the Wellcome Trust and Royal Society through a Sir Henry Dale Fellowship to JKB (98418/Z/12/Z).

